# Microgliosis, astrogliosis and loss of aquaporin-4 polarity in frontal cortex of COVID-19 patients

**DOI:** 10.1101/2024.04.10.588851

**Authors:** Antonia Beiersdorfer, Natalie Rotermund, Kristina Schulz, Miriam Busch, Daniela Hirnet, Stephan Henne, Florian L. Schwarzenberg, Matthias Dottermusch, Benjamin Ondruschka, Jakob Matschke, Clemens Wülfing, Markus Glatzel, Christian Lohr

**Author notes:** Equal contribution (Antonia Beiersdorfer, Natalie Rotermund, Kristina Schulz). **Data availibility statement:** All data of this study will be made available by the authors upon request. **Ethics approval statement:** The study adhered to the Declaration of Helsinki principles, and the Ethics Committee of the Hamburg Chamber of Physicians was informed (PV7311 and 2020-10353-BO-ff).

## Abstract

The severe acute respiratory syndrome coronavirus type 2 (SARS-CoV-2), causing human coronavirus disease 2019 (COVID-19), not only affects the respiratory tract, but also impacts other organs including the brain. A considerable number of COVID-19 patients develop neuropsychiatric symptoms that may linger for weeks and months and contribute to “long-COVID”. While the neurological symptoms of COVID-19 are well described, the cellular mechanisms of neurologic disorders attributed to the infection are still enigmatic. Here, we studied the effect of an infection with SARS-CoV-2 on the structure and expression of marker proteins of astrocytes and microglial cells in the frontal cortex of patients who died from COVID-19 in comparison to non-COVID-19 controls. Most of COVID-19 patients had microglial cells with retracted processes and rounded and enlarged cell bodies in both gray and white matter, as visualized by anti-Iba1 staining and confocal fluorescence microscopy. In addition, gray matter astrocytes in COVID-19 patients were frequently labeled by intense anti-GFAP staining, whereas in non-COVID-19 controls, most gray matter astrocytes expressed little GFAP. The most striking difference between astrocytes in COVID-19 patients and controls was found by anti-aquaporin-4 (AQP4) staining. In COVID-19 patients, a large number of gray matter astrocytes showed an increase in AQP4. In addition, AQP4 polarity was lost and AQP4 covered the entire cell, including the cell body and all cell processes, while in controls, AQP4 immunostaining was mainly detected in endfeet around blood vessels and did not visualize the cell body. In summary, our data suggest neuroinflammation upon SARS-CoV-2 infection including microgliosis and astrogliosis, including loss of AQP4 polarity.

## 1 INTRODUCTION

Neuroinflammation as evoked by traumatic brain injury, autoimmune responses or infection results in activation of microglial cells and astrocytes (das Neves, Sousa, Sousa, Cerqueira, & Marques, 2021; Linnerbauer, Wheeler, & Quintana, 2020; Patani, Hardingham, & Liddelow, 2023). Infection with severe acute respiratory syndrome-corona virus type 2 (SARS-CoV-2) causes human coronavirus disease 2019 (COVID-19) which is frequently associated with neuroinflammatory responses (Almutairi, Sivandzade, Albekairi, Alqahtani, & Cucullo, 2021; Gonçalves de Andrade et al., 2021; Iadecola, Anrather, & Kamel, 2020; Matschke et al., 2022; Matschke et al., 2020; A. C. Yang et al., 2021). Although being mainly a respiratory disease, COVID-19 may lead to severe neurological disorders such as fatigue, headaches, memory and mood impairment (Bodnar, Patel, Ho, Luo, & Hu, 2021; Gonzalez-Fernandez & Huang, 2023; Hensley, Markantone, & Prescott, 2022; Penninx, Benros, Klein, & Vinkers, 2022). Both neurological as well as non-neurological symptoms persist after subsidence of the infection in a considerable number of patients, a syndrome known as post acute sequelae of COVID-19 (PASC), or simply post-COVID or long-COVID (Goldstein, 2024; Nalbandian et al., 2021; Song et al., 2021; Steardo, Steardo, Verkhratsky, & Scuderi, 2021). Recent investigations found an increased risk of incident neurological sequelae such as stroke, migraine, seizures, mental health disorders, Guillain-Barre syndrome and encephalitis in long-COVID patients and late effects of COVID-19 may become a major health issue in the future (Aghajani Mir, 2023; Brown et al., 2023). Therefore, understanding the molecular and cellular consequences of the viral infection within the central nervous system is inevitable to develop treatment of long-COVID.

Several publications report invasion of SARS-CoV-2 into brain cells such as neurons and glial cells, but whether the number of virus particles found in brain tissue is large enough to explain severe neurological manifestations is still under debate (Y. Chen, Yang, Chen, & Cui, 2022; Cosentino et al., 2021; Krasemann, Dittmayer, et al., 2022; Meinhardt et al., 2024; Monje & Iwasaki, 2022; Solomon, 2021; Stein et al., 2022). Alternatively, neuroinflammation triggered by the “cytokine storm” originating from infected epithelial cells of respiratory organs may cause those mental and neurological disorders as described above (Gupta et al., 2020; Liu et al., 2020; Monje & Iwasaki, 2022; Wu & Tang, 2020). Post-mortem studies of COVID-19 brain tissue focused on immune cells and neurons, while only few studies addressed astrocytes (Tremblay, Madore, Bordeleau, Tian, & Verkhratsky, 2020; Vargas et al., 2020). Initially described as support cells with homeostatic function only, astrocytes have been found to be directly involved in neuronal information processing such as synaptic transmission and plasticity (Santello, Toni, & Volterra, 2019; Semyanov, Henneberger, & Agarwal, 2020). In addition, astrocytes are affected by and are often causal of neurological diseases (Lee, Wheeler, & Quintana, 2022; Lohr, 2023; Patel, Tewari, Chaunsali, & Sontheimer, 2019). In COVID-19, reactive astrogliosis has been reported, including increased expression of glial fibrillary acidic protein (GFAP), and glial markers were found elevated in plasma and cerebrospinal fluid (Hanson et al., 2022; Huang & Fishell, 2022; Kanberg et al., 2020; Rosu et al., 2022; Spanos et al., 2022; Steardo, Steardo, & Scuderi, 2022; Tremblay et al., 2020). In the present study, we investigated the astrocyte markers GFAP, calcium-binding protein S100B and aquaporin-4 (AQP4) as well as microgliosis in frontal lobe of COVID-19 brains using immunohistochemical staining and compared the results with those from control brains. We found microgliosis as well as astrogliosis and loss of AQP4 polarity in most of COVID-19 patients, while in the control group, only few samples showed signs of microgliosis and astrogliosis. Our results suggest that COVID-19 results in severe neuroinflammation and activation of both microglia and astroglia.

## 2 METHODS

### 2.1 Autopsies and ethical considerations

In the state of Hamburg, Germany, numerous individuals who died from or with SARS-CoV-2 infection between March 1st and December 31st 2020 underwent full autopsies at the Institute of Legal Medicine, University Medical Center Hamburg-Eppendorf (UKE) (Edler et al., 2020; Fitzek et al., 2021). SARS-CoV-2 infection was diagnosed via throat swabs followed by immediate RT-qPCR for viral RNA. Clinical data, including medical history and ante mortem findings, were collected. During autopsies, the whole undissected brain with the dura mater and the pituitary were kept in formalin for at least two weeks, followed by full neuropathological assessment in the Institute of Neuropathology of the UKE. The study adhered to the Declaration of Helsinki principles, and the Ethics Committee of the Hamburg Chamber of Physicians was informed (PV7311 and 2020-10353-BO-ff).

Age-matched historical controls predating 2019 were selected from the Institute of Neuropathology’s archives. Exclusion criteria included pre-existing neurological diseases, sepsis, and intensive care-related interventions. The study encompassed frontal sections and tissue samples extracted from the frontal cortex (FC) from twenty-two individuals, categorized into two distinct groups, twelve individuals diagnosed with COVID-19 and ten control subjects (Suppl. table 1).

### 2.2 Immunohistochemistry

Standard histopathological stainings were performed on 8 μm thick formalin-fixed paraffin-embedded sections. For deparaffinization, slides were immersed in Roti-Histol (Carl Roth, Karlsruhe, Germany) for two cycles of 5 minutes each, followed by a descending series of ethanol using 100 % (5 min), 95 % (5 min), 70 % (10 min), and 50 % (5 min). Subsequently, sections were rinsed with deionized water for 5 minutes. Antigen retrieval varied depending on the primary antibodies (Suppl. table 2). Alkaline antigen retrieval (10 mM Tris; 1 mM EDTA; 0,05% Tween-20, pH 9, 95°C for 10 min) was used for staining with anti-CD31 antibody, while acidic antigen retrieval (10 mM citric acid, 0.05% Tween-20, pH 6.0, 95°C for 10 min) was used for all other stainings. After heat treatment, sections were allowed to cool at room temperature for 20 minutes. Prior to primary antibody incubation, sections were treated with PBST (phosphate-buffered saline with Tween-20) for 60 minutes on a shaker at room temperature. Subsequently, sections were incubated in blocking solution (5 % BSA, 0.4 % Triton-X100) for 60 minutes at room temperature in a humid chamber. Slices were then incubated overnight in primary antibody solution (5 % BSA, 0.2 % Triton-X100, 0.1 % sodium acid in PBS) with antibodies listed in supplementary table 2 at 4°C. To quench autofluorescence, Biotium TrueBlack (Biozol, Echingen, Germany) diluted 1:20 in 70 % ethanol was added to the brain sections for 30 seconds. Specimens were then incubated with the secondary antibody solution for 1 hour at room temperature.

### 2.3 Immunohistochemistry of free-floating frontal cortex slices

To collect three-dimensional image stacks of astrocytes, paraffin-embedded tissue blocks were sectioned 150 μm thick using a vibratome (VT1000S, Leica, Germany). Brain sections were heated to a temperature of approximately 60°C on a hotplate until the wax phase transformed into a liquid state. Subsequently, the sections were deparaffinized in Roti-Histol at 36°C in two cycles. Brain sections were then immersed in descending concentrations of ethanol; 100 % (10 min), 95 % (10 min), 70 % (20 min), and 50 % (10 min), followed by immersion in deionized water (5 min). The sections were subjected to antigen retrieval by immersing them in an acidic buffer at 95°C for 10 minutes and subsequently cooling to room temperature for 20 minutes. The tissue was permeabilized with PBST for 90 minutes and incubated in blocking solution (5 % BSA, 0.4 % Triton-X100) for 120 minutes. Anti-AQP4 antibody in antibody solution (5 % BSA, 0.2 % Triton-X100, 0.1 % sodium acid in PBS) was added, dishes were sealed and incubated at 4°C for a period of 6 days. To reduce autofluorescence, specimens were incubated in Biotium TrueBlack (1:20 in 70 % EtOH) for 2 minutes and subsequently incubated in secondary antibody solution with 0.1 % sodium azide in PBS protected from light at 4°C for 6 days. To optimize tissue transparency for 3D imaging, sections were transferred to RapiClear 1.49 (SunJin Lab, Hsinchu City, Taiwan) for a period of 60 minutes. Sections were then mounted on slides using spacers, a cover slip and RapiClear as mounting medium.

### 2.4 Confocal microscopy, image processing, and quantitative image analysis of AQP4 distribution

Single plane images were acquired with a confocal microscope (eC1, Nikon, Düsseldorf, Germany) using 10x/NA 0.5 and 40x/NA 1.3 objectives. Z-stacks of images from cleared brain slices were imaged with the 40x/NA 1.3 objective with 150 nm axial spacing. Image stacks were deconvolved using Huygens software (SVI, Hilversum, Netherlands). Both, single plane images and projections from image stacks were adjusted for brightness and contrast with Adobe Photoshop (Dublin, Ireland). In addition, for a quantitative analysis of AQP4 distribution, high resolution large area stitches from 8 μm thick specimens were captured at high magnifications using an Axio Imager.M2m microscope (Zeiss, Oberkochen, Germany). An average of 950 images per brain slice were acquired using a 20x/0.8 objective. Regions of interest, encompassing both gray and white matter, were delineated within the ZEN software’s tile region configuration (Zeiss). A 3x3 grid of focal points was generated for each image area, with manual focus adjustments made as necessary. Images were systematically acquired across preselected regions and seamlessly amalgamated into comprehensive stitched images. This process facilitated a holistic understanding of AQP4 distribution patterns. Analysis of AQP4 distance to capillaries was performed on high resolution large area stitches using Imaris 9.7.2 (Oxford Instr., UK). “Hot spots” of AQP4 immunostaining were defined automatically by identifying local maxima of fluorescence and the distance of “hot spots” to the nearest blood vessel, stained with anti-CD31 antibody, was measured.

### 2.5 Statistical analysis

Data analysis was conduced using OriginPro 2021 (OriginLab Corporation, Northampton, MA, USA). The Shapiro-Wilk test was used to test the Gaussian distribution of the data. For comparison between case-control populations the Student’s t-test was used. Cumulative distribution of distances between AQP4 “hot spots” and blood vessels were tested for significant differences between COVID-19 and controls using the two-sample Kolmogorov-Smirnov test. *P*-values of 0.05 or less were considered to be statistically significant.

## 3 RESULTS

We studied brain tissue from twelve COVID-19 individuals (8 male, 4 female) and ten deceased as control (7 male, 3 female). Average age was 69.0 +/- 19.6 years (range 29-93 years) for COVID-19 cases and 67.9 +/- 17.1 (range 33-92 years) for controls. The difference in age was not significantly different (*P*=0.899). Average brain weights were 1360 +/- 52.8 g for COVID-19 cases and 1330 +/- 63.8 g for controls, the difference not being significant. Neither COVID-19 patients nor controls showed any significant macroscopic damage to the brain, such as cerebral infarction, bleeding, tumor, trauma, or microscopic findings, such as demyelination, meningitis, encephalitis, endotheliitis, or widespread hyaline microthrombosis. As the result of autopsy, all COVID-19 cases died from COVID-19. For age and sex details of cases and controls included see Suppl. table 1.

### 3.1 Microgliosis in frontal lobes of COVID-19 patients

We studied the morphology of microglial cells by fluorescent immunostaining using antibodies against Iba1, a calcium binding protein expressed by macrophages including microglial cells. In both gray and white matter of control patients, microglial cells possessed long, ramified processes, indicative of homeostatic microglia (Fig. 1a-d). In COVID-19 patients, in contrast, microglial cells partly retracted their processes and had the appearance of reactive microglial cells (Fig. 1e-h). Overall, 11 out of 12 COVID-19 patients showed signs of reactive microgliosis, while microglial cells in only 1 out of 10 control patients were reactive. This indicates severe neuroinflammation in frontal lobes of patients that died of COVID-19. We co-stained endothelial cells with anti-CD31 antibodies to highlight blood vessels and to assess whether perivascular microglial cells were more affected by COVID-19 compared to parenchymatous microglial cells. We could not detect any difference in the reactive stage of microglial cells regarding of their relation to blood vessels.

**Figure 1.**
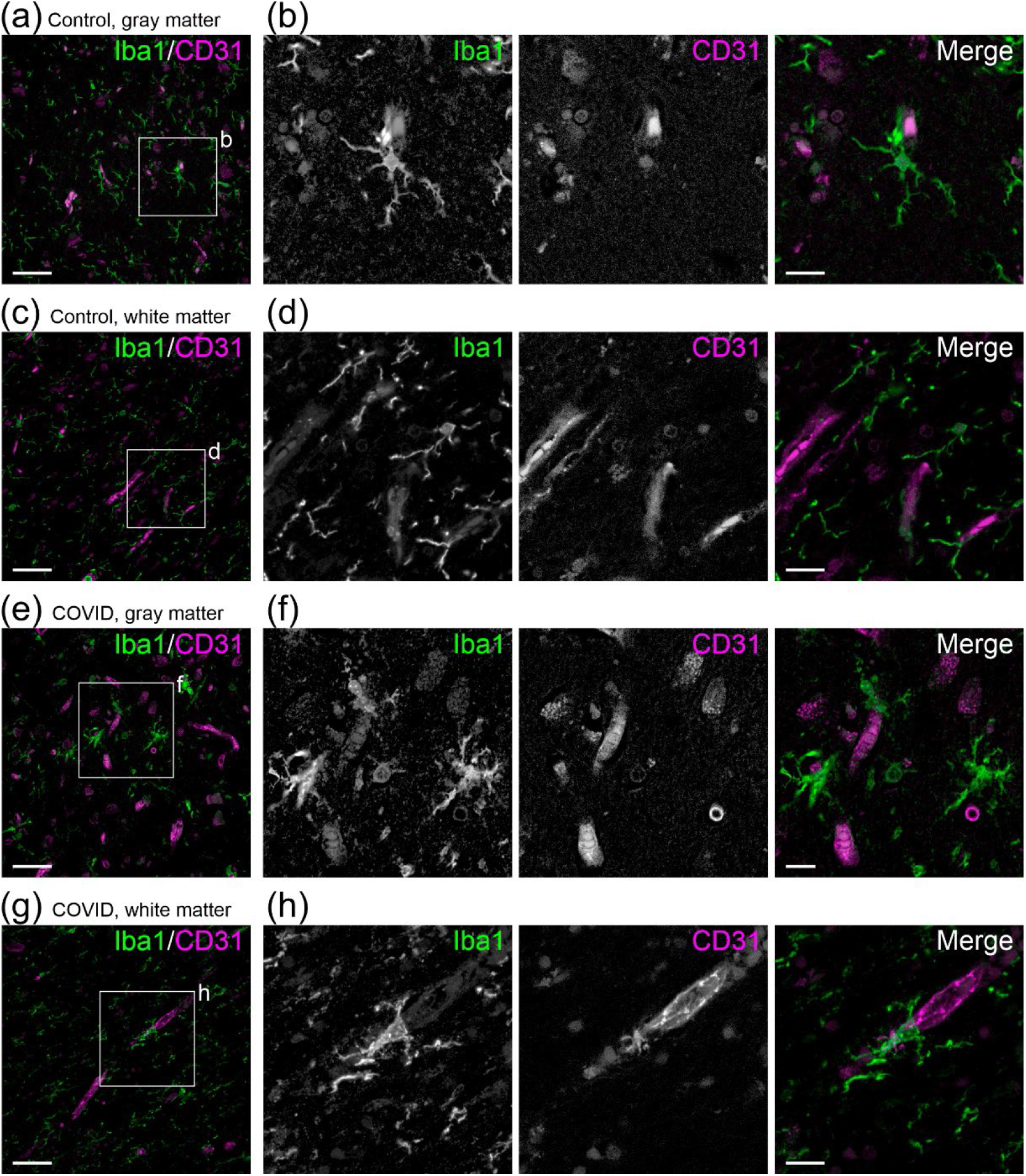
Reactive microgliosis in frontal lobe of COVID-19 patients. (a) Microglial cells in gray matter labeled with anti-Iba1 antibody (green) and endothelial cells labeled with anti-CD31 antibody (magenta). (b) Magnified section of (a). Microglial cells in gray matter of control brain tissue possessed small cell bodies and ramified processes, typical for homeostatic microglial cells. (c) Anti-Iba1 and anti-CD31 staining of white matter in control patients. (d) Homeostatic microglial cells in white matter of control patients. (e) Anti-Iba1 and anti-CD31 staining in gray matter brain tissue of COVID-19 patients. (f) Magnified section of (e). Processes of microglial cells were shortened and thickened and the cell bodies were bulky, indicating reactive microgliosis. (g) Microglial cells in white matter of COVID-19 patients. (h) Reactive microglial cells in white matter of COVID-19 patients. Scale bars in (a), (c), (e), (g): 50 μm; in (b), (d), (f), (h): 15 μm.

### 3.2 Increased GFAP expression in gray matter astrocytes of COVID-19 patients

We stained for GFAP to assess the effect of COVID-19 on GFAP expression in frontal cortex and white matter. We found GFAP-positive astrocytes in white matter in both control and COVID-19 tissue, while in gray matter astrocytes intense GFAP staining was found only in COVID-19 patients in larger numbers (Fig. 2a, c). We further stained S100B to highlight all astrocytes regardless of their GFAP expression and compared GFAP immunoreactivity with S100B immunoreactivity in cortical gray matter at higher magnification. In controls, astrocytes labeled by S100B antibodies mostly expressed little GFAP (Fig. 2b). In COVID-19 patients, in contrast, a large fraction of S100B-positive gray matter astrocytes was co-labeled by GFAP, whereas some S100B-positive astrocytes were GFAP-negative (Fig. 2d). Ten out of twelve COVID-19 patients (83%) had significant expression of GFAP in gray matter astrocytes in the frontal cortex, but in only four out of ten controls (40%), astrocytes were clearly GFAP-positive. In white matter, astrocytes highly expressed GFAP in both controls and COVID-19, and no difference in GFAP expression between controls and COVID-19 patients was found (Fig. 3).

**Figure 2.**
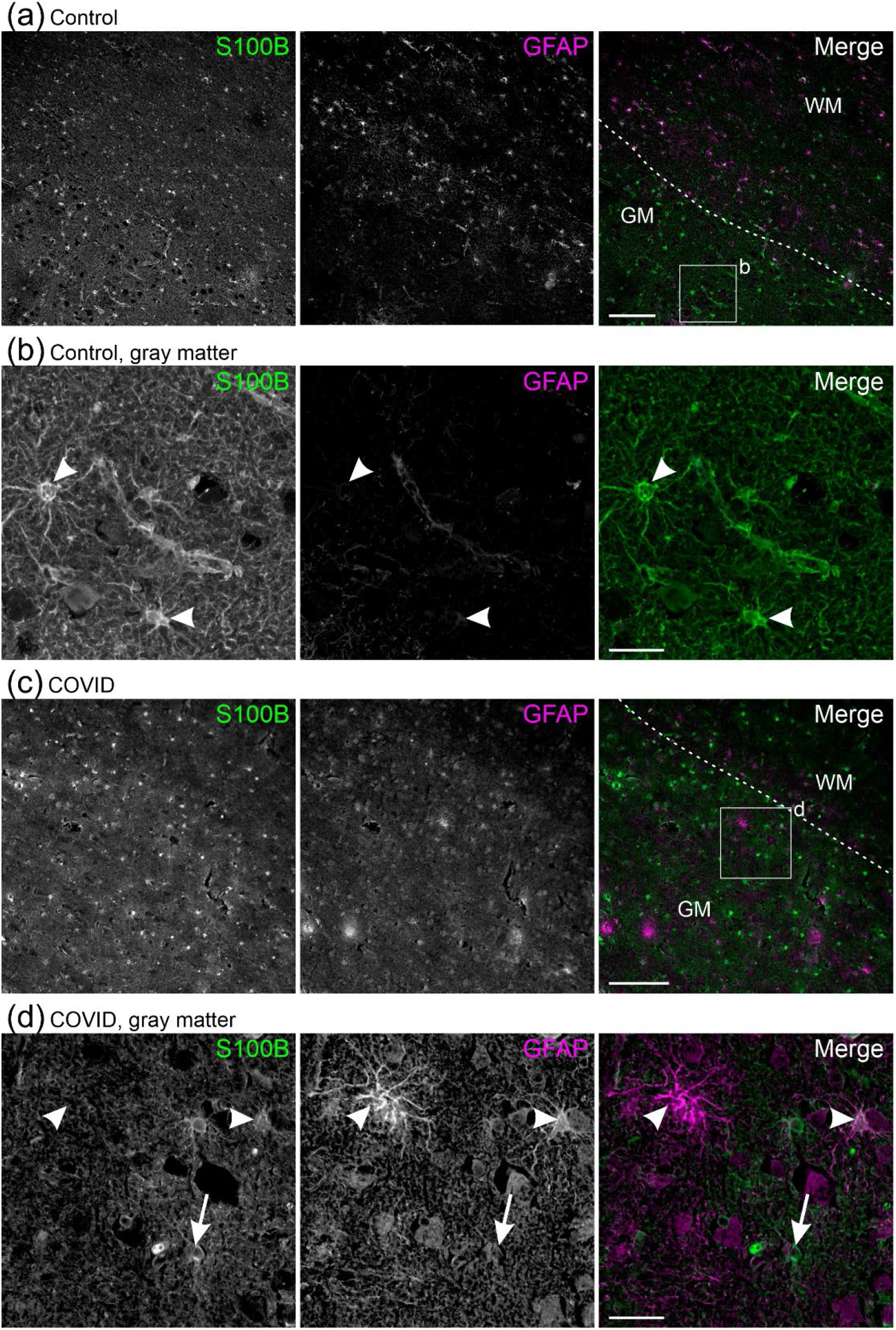
Increased GFAP expression in COVID-19 patients. (a) Anti-S100B and anti-GFAP co-staining in gray matter (GM) and white matter (WM) frontal cortex of control patients. (b) S100B-positive gray matter astrocytes (arrowheads) were GFAP-negative. (c) Anti-S100B and anti-GFAP co-staining in frontal cortex of COVID-19 patients. (d) In COVID-19, some gray matter astrocytes were GFAP-positive (arrowheads), while others lacked GFAP (arrow). Scale bars in (a) and (c): 200 μm; in (b) and (d): 30 μm.

**Figure 3.**
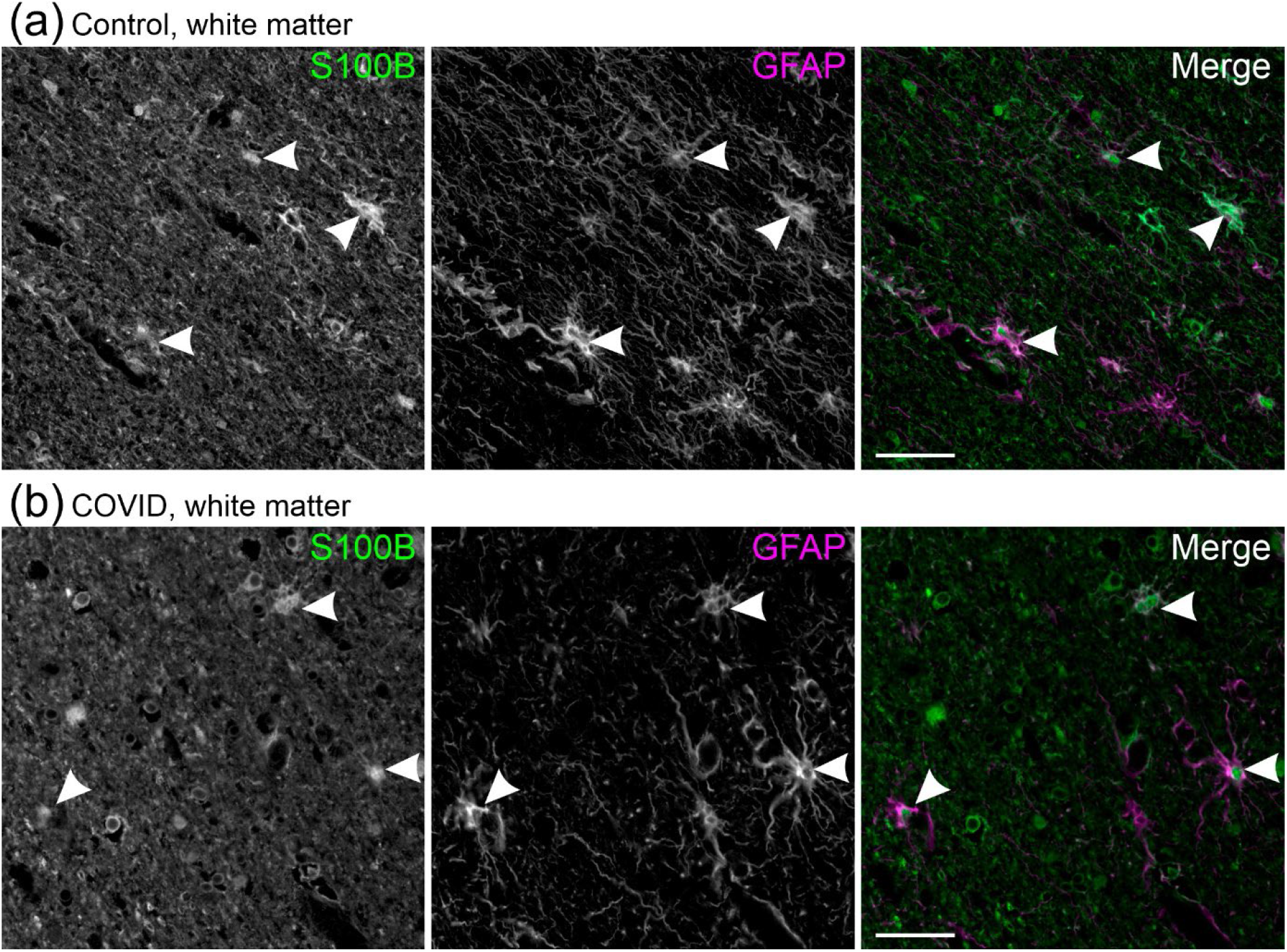
GFAP expression in white matter astrocytes. (a) Anti-S100B and anti-GFAP co-staining in white matter of control patients and (b) COVID-19 patients revealed GFAP expression in most of S100B-expressing astrocytes (arrowheads). Scale bars: 30 μm.

### 3.3 GFAP expression is increased in parenchymatous and perivascular gray matter astrocytes in COVID-19 patients

Astrocytes ensheath blood vessels in the brain by a layer of so-called endfeet. Hence, cytokines as well as immune cells invading the central nervous system parenchyma from the bloodstream will impact perivascular astrocytes earlier and stronger than parenchymatous astrocytes. We studied the relation of GFAP-positive astrocytes to blood vessels, labeled by antibodies against the endothelial marker CD31, to assess whether perivascular astrocytes show a stronger expression of GFAP than parenchymatous astrocytes. As shown before, gray matter astrocytes of controls were weakly stained for GFAP, whereas white matter astrocytes of controls as well as gray and white matter astrocytes of COVID-19 patients were intensely labeled by GFAP antibodies (Fig. 4a-d). In later categories, parenchymatous astrocytes were found to be GFAP-positive. In addition, GFAP-positive sheaths around blood vessels were visible, indicating increased GFAP levels in the endfeet.

**Figure 4.**
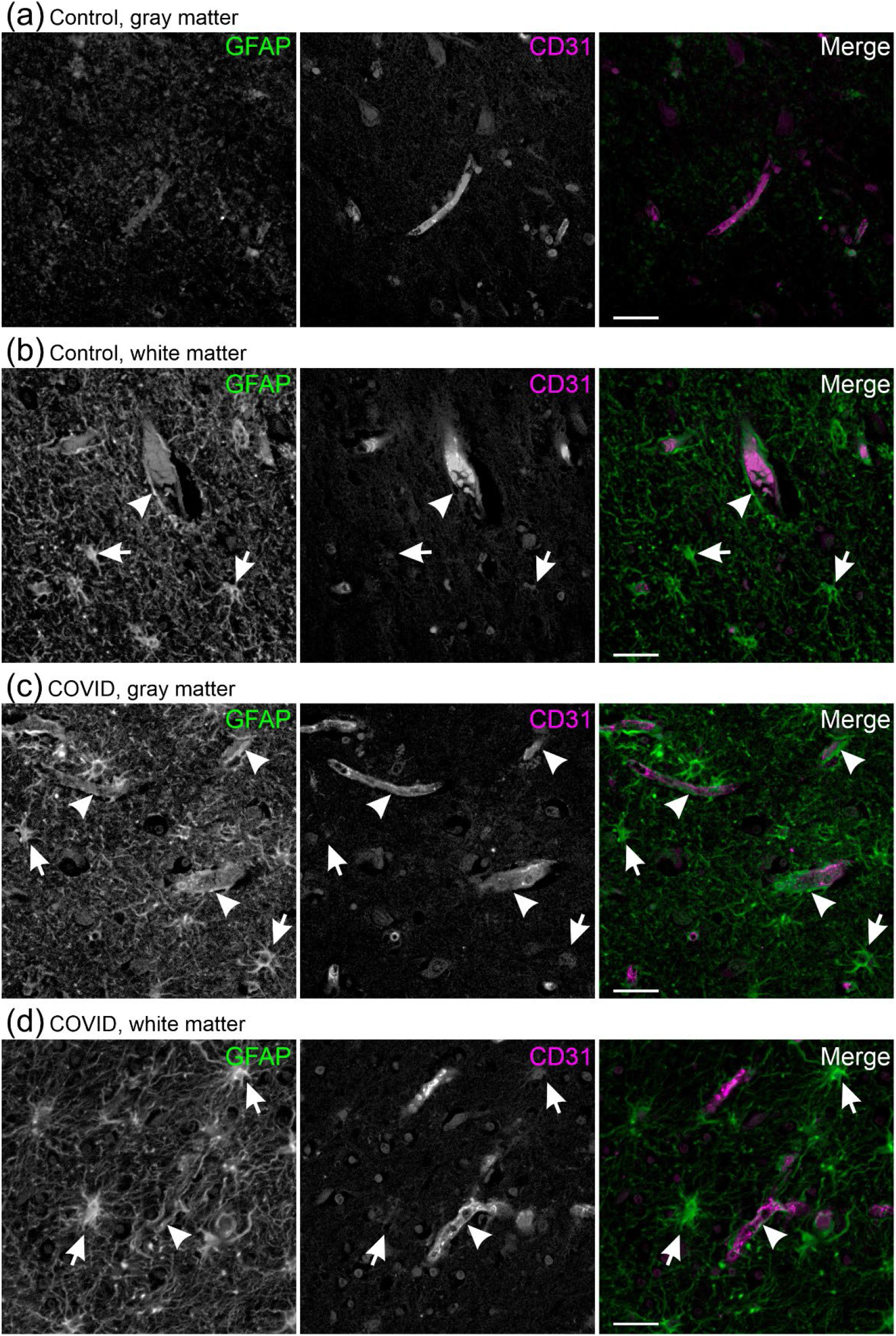
GFAP is increased in COVID-19 patients in perivascular and parenchymatous astrocytes. (a) Low GFAP expression in gray matter of control patients. (b) In white matter of control patients, parenchymatous astrocytes (arrows) and perivascular endfeet (arrowheads) were GFAP positive. (c) In COVID-19 patients, parenchymatous astrocytes and perivascular endfeet in gray matter and (d) white matter were GFAP-positive. Scale bars: 50 μm.

### 3.4 Increased AQP4 expression in COVID-19 patients

We were interested in whether an infection with SARS-CoV-2 affects the expression of AQP4 and compared cortical tissue from COVID-19 patients with that of controls. Anti-AQP4 staining revealed a low expression in control patients (Fig. 5a). In COVID-19 patients, in contrast, AQP4 expression was increased, which was most obvious in gray matter (Fig. 5b). Increased AQP4 immunoreactivity was found in twelve out of twelve COVID-19 patients (100%) and a moderate increase in AQP4 immunoreactivity in three out of ten control patients (30%), while the remaining control patients had weak AQP4 expression. At higher magnification, single strongly AQP4-positive astrocytes were found only sporadically in most control patients, whereas in COVID-19 patients AQP4-positive astrocytes were detected at high density (Fig. 5c-f). To reveal a better impression of the distribution of AQP4 in individual astrocytes of COVID-19 patients, we prepared tissue slices of 300 μm thickness, cleared the tissue and imaged astrocytes at high resolution, resulting in confocal image stacks that covered the entire cell in three dimensions. After deconvolution of the image stacks, the data set was rendered. Figure 5g shows a projection of a subset of images of the image stack, covering the cell body and some processes of an astrocyte with high AQP4 expression. Strong AQP4 immunoreactivity was found in the entire cell, including the cell body. Our data show that neuroinflammation due to COVID-19 results in strong upregulation of AQP4 in a large number of cortical astrocytes.

**Figure 5.**
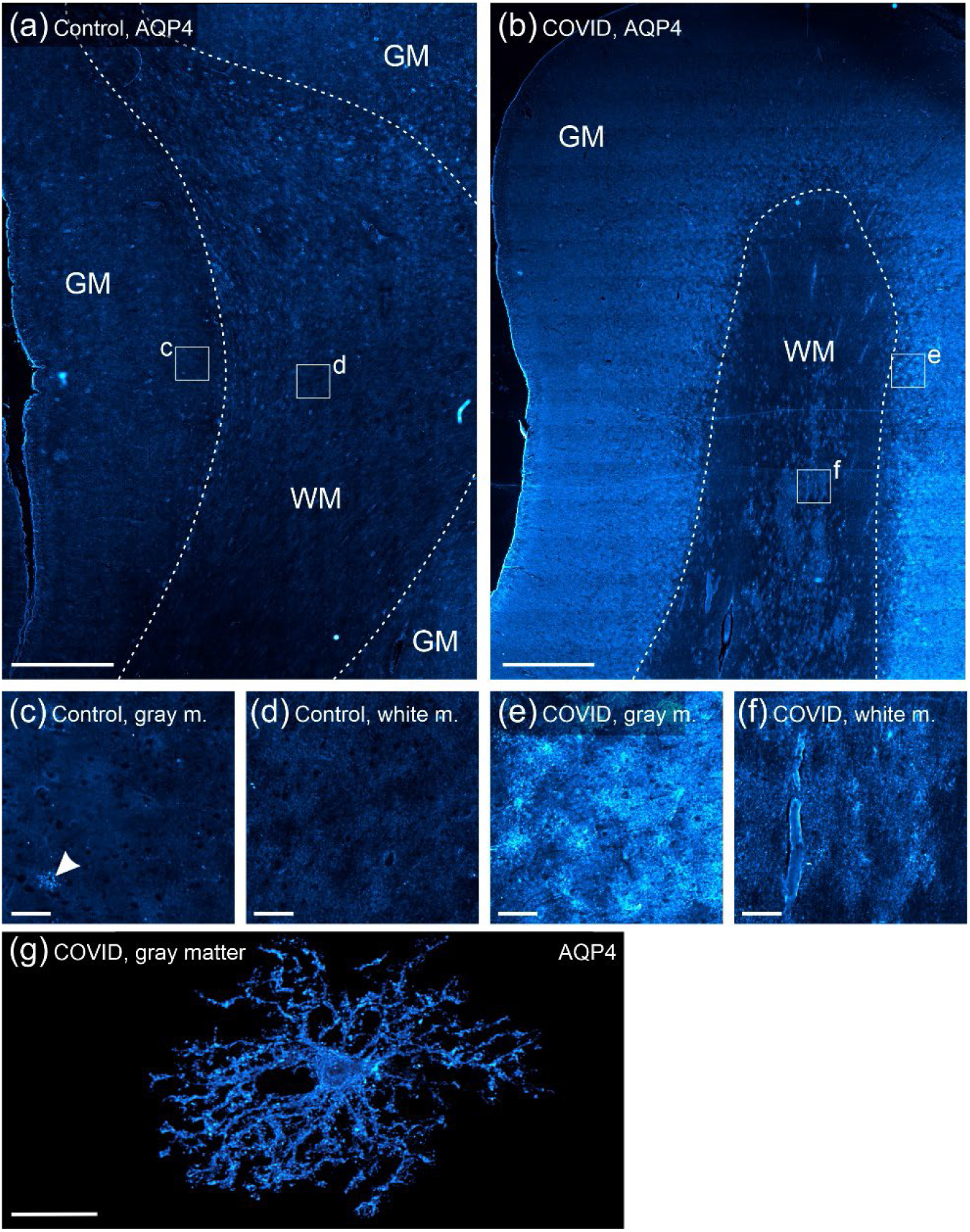
Increased AQP4 expression in COVID-19 patients. (a) AQP4 antibody staining of frontal cortex of control patients and (b) COVID-19 patients. (c) AQP4 immunoreactivity of control patients at higher magnification in gray matter and (d) white matter. (e) AQP4 immunostaining was increased in gray matter and (f) white matter of COVID-19 patients. (g) Projection of a three-dimensional stack of confocal images of AQP4 staining in cleared tissue. The image stack was post-hoc processed by deconvolution and one astrocyte was masked by manually outlining cell structures and removing background fluorescence. Scale bars: 2 mm in (a) and (b); 100 μm in (c)-(f); 25 μm in (g).

### 3.5 Loss of AQP4 polarity in COVID-19 patients

Both AQP4 and GFAP immunoreactivity was low in gray matter and AQP4 was most obvious around blood vessels in controls (Fig. 6a). In COVID-19, a large number of astrocytes was strongly AQP4- and GFAP-positive (Fig. 6b). Some of AQP4-positive astrocytes were also GFAP-positive, however, there were strongly GFAP-positive astrocytes that possessed no or only weak AQP4 immunoreactivity in the cell body and large cell processes. This indicates that in COVID-19, astrogliosis that leads to increased GFAP expression does not necessarily result in increased AQP4 expression in the same cell. We next assessed the polarized localization of AQP4 by analyzing the distance of AQP4 “hot spots” to the nearest blood vessel labeled by anti-CD31 (Fig. 6c, d). The cumulative distribution plot of the distance of AQP4 immunostaining and blood vessels revealed a significantly closer localization of AQP4 to blood vessels in controls compared to COVID-19 patients (Kolmogorov-Smirnov, p<0.001), suggesting loss of polarity of AQP4 distribution (Fig. 6e).

**Figure 6.**
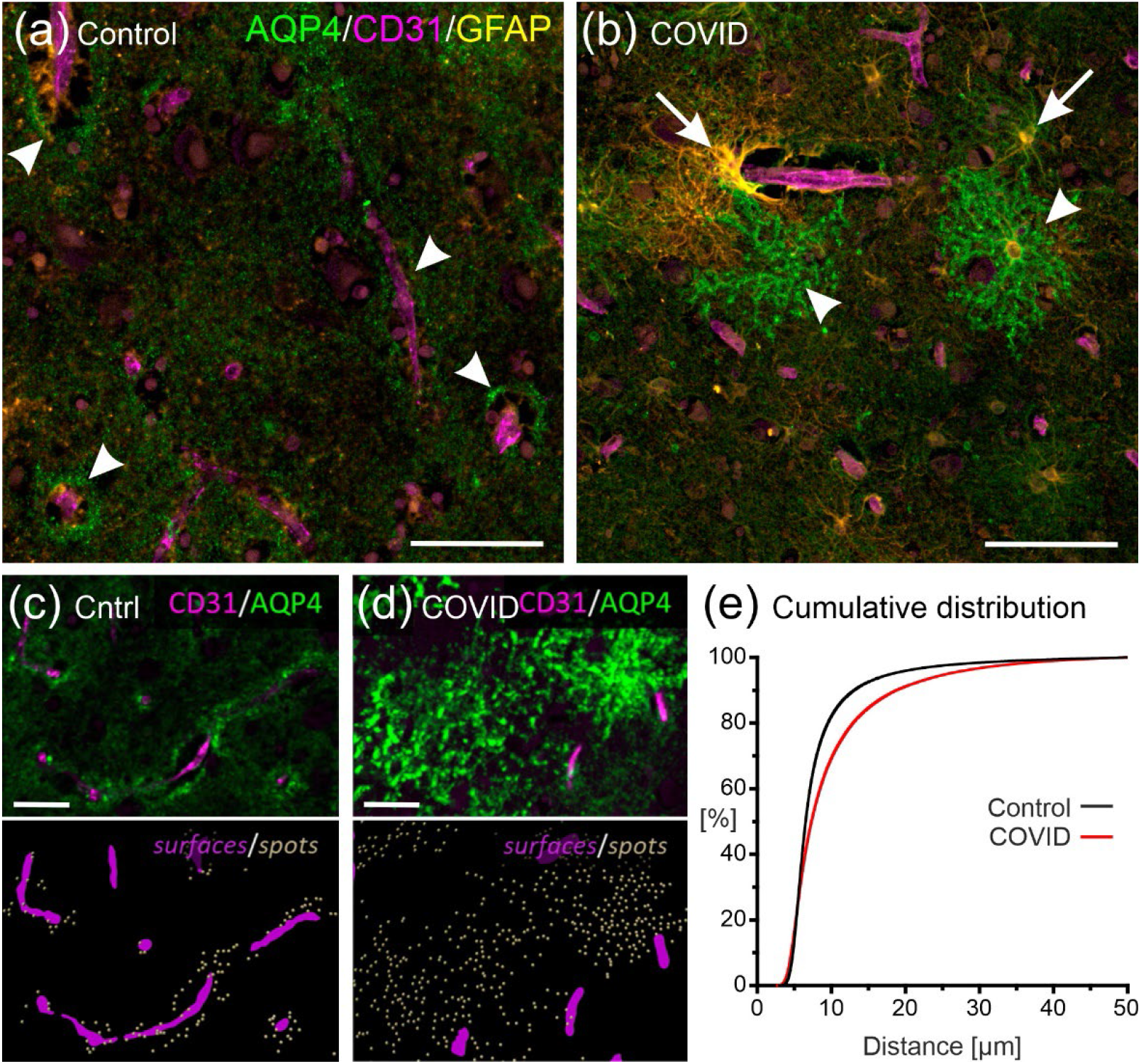
Loss of AQP4 polarity in COVID-19 patients. (a) Triple staining of AQP4 (green), GFAP (yellow) and CD31 (magenta) revealed low expression of AQP4 and GFAP in gray matter of controls, with perivascular localization of AQP4 (arrowheads). (b) In gray matter of COVID-19 cases, high expression AQP4 and GFAP was found. High expression of AQP4 and GFAP colocalized in some astrocytes (arrowheads), while some strongly GFAP-positive astrocytes possessed no or low AQP4 immunoreactivity (arrows). (c) CD31-labeled blood vessels (magenta) and AQP4 were stained (upper panel), the blood vessels were segmented and hot spots of AQP4 immunoreactivity were visualized to measure the distance of AQP4 to the closest blood vessel in controls and (d) COVID-19 cases. (e) The cumulative distribution plot shows that in COVID-19 cases the distance of AQP4 to blood vessels was larger compared to controls (p<0.001), indicating loss of AQP4 polarity. Scale bars: 50 μm in (a) and (b); 20 μm in (b) and (d).

## 4 DISCUSSION

COVID-19 is known to cause severe neurological and psychological symptoms but the role of astrocytes in COVID-19-related brain disorders has been studied only insufficiently. In the present study, we have investigated the expression and subcellular distribution of glial markers in post mortem frontal cortex tissue of COVID-19 patients by immunofluorescence. We found reactive microgliosis, as indicated by rounded Iba1-positive microglial cells with shortened cell processes, and astrogliosis indicated by increased GFAP expression in gray matter compared to control patients, implying neuroinflammation in COVID-19. In addition, in COVID-19 patients a large number of astrocytes possessed strong AQP4 immunoreactivity and homogeneous distribution of AQP4 throughout the entire cell, while AQP4 in control patients was accumulated in astrocyte endfeet around blood vessels.

### SARS-CoV-2 causes neuroinflammation

Several studies reported signs of neuroinflammation in tissue from COVID-19 patients (reviewed by (Almutairi et al., 2021; Brown et al., 2023; Gonçalves de Andrade et al., 2021; Hernández-Parra et al., 2023; Iadecola et al., 2020; Natale, Lukens, & Petri, 2022; Tremblay et al., 2020; Vanderheiden & Klein, 2022; A. C. Yang et al., 2021)). SARS-CoV-2 not only infects the respiratory tract but also other organs, including the brain and its cerebrovasculature (Y. Chen et al., 2022; Jakhmola, Indari, Chatterjee, & Jha, 2020; Krasemann, Dittmayer, et al., 2022; Monje & Iwasaki, 2022; Solomon, 2021). Virus particles might invade into the brain through the olfactory nerve or through the choroid plexus and cause neuroinflammation (A. C. Yang et al., 2021). Angiotensin converting enzyme 2 (ACE2), the key receptor for SARS-CoV-2 entry into host cells, is highest expressed in the brain by astrocytes, although still at low levels as compared to other organs such as lungs, kidneys and heart (Li, Hasson, Daggumati, Zhang, & Thorek, 2023). In theory, thus, astrocytes could be the main target of the virus particles in the brain, causing astrogliosis and neuroinflammation. However, the number of virus particles in brain cells including astrocytes found by most studies is too low to explain severe neurological symptoms (R. Chen et al., 2020; Cosentino et al., 2021; Krasemann, Dittmayer, et al., 2022; Monje & Iwasaki, 2022; Solomon, 2021). More likely, neuroinflammation is the result of the extreme high concentration of pro-inflammatory cytokines released by infected cells of the respiratory tract, reaching the brain via the bloodstream and causing impairment of the blood-brain barrier integrity with subsequent invasion of immune cells into the brain parenchyma (Gupta et al., 2020; Krasemann, Haferkamp, et al., 2022; Liu et al., 2020; Meinhardt et al., 2024; Monje & Iwasaki, 2022; Shabani, Liu, & Su, 2023). Among the cytokines are interleukin 2R, interleukin 6, interleukin 8, tumor necrosis factor (TNF, also known as TNF-alpha) and, interestingly, also the anti-inflammatory interleukin-10 (Devlin & Gombolay, 2023; Liu et al., 2020). Macrophages and lymphocytes penetrate the blood-brain barrier, activate microglial cells and astrocytes and finally determine neuroinflammation (Almutairi et al., 2021; Brown et al., 2023; Gonçalves de Andrade et al., 2021; Hernández-Parra et al., 2023; Iadecola et al., 2020; Tremblay et al., 2020; Vanderheiden & Klein, 2022). In our study, both microglial cells and astrocytes showed an activated phenotype in COVID-19 patients, confirming neuroinflammation. We found microgliosis and astrogliosis not only in perivascular cells, but in the entire parenchyma. This suggests an advanced stage of inflammation in which pro-inflammatory factors and immune cells migrate from the direct environment of the vasculature into the parenchyma.

### Loss of AQP4 polarity in COVID-19 patients

In addition to GFAP, one of the most often used markers for astrogliosis, we employed an antibody against AQP4 to study astrocytes in COVID-19. In healthy astrocytes, subcellular AQP4 distribution is polarized and AQP4 is mainly located in the astrocyte endfeet surrounding blood vessels, where it functionally contributes to the blood-brain barrier and is involved in extracellular volume dynamics, water homeostasis and waste clearance (Nagelhus & Ottersen, 2013; Rasmussen, Mestre, & Nedergaard, 2022; Verkman, Smith, Phuan, Tradtrantip, & Anderson, 2017). We found increased AQP4 expression and loss of AQP4 polarization in all COVID-19 patients investigated, in contrast to a former study that showed decreased AQP4 (Rosu et al., 2022). Malfunction of AQP4 contributes to the progression of neurodegenerative diseases such as Alzheimer’s and Parkinson’s disease, and AQP4 autoantibodies cause neuromyelitis optica spectrum disorder (Buccellato, D’Anca, Serpente, Arighi, & Galimberti, 2022; Mader & Brimberg, 2019; Nedergaard & Goldman, 2020; Pittock, Zekeridou, & Weinshenker, 2021). Increased expression and depolarized subcellular distribution of AQP4 has been shown in animal models of neuroinflammation and neurodegeneration such as traumatic brain injury (Ren et al., 2013), local ischemia (Wang et al., 2012), edema (Tourdias et al., 2011), prion disease (Kushwaha et al., 2023) and Alzheimer’s disease (Feng et al., 2023; Pedersen, Keil, Han, Wang, & Iliff, 2023; J. Yang et al., 2011). Similar phenotypes were observed in human brain tissue in subjects with Alzheimer’s disease (Hoshi et al., 2012; Valenza, Facchinetti, Steardo, & Scuderi, 2019; Zeppenfeld et al., 2017), Creitzfeldt-Jakob disease (Rodríguez et al., 2006) and stroke (Roşu et al., 2019). Impairment of ion and water homeostasis as well as waste clearance due to loss of AQP4 polarization exacerbate the neurological outcome of these disorders or even is causal of them (Mader & Brimberg, 2019; Nedergaard & Goldman, 2020; Pittock et al., 2021; Verkman, 2012; Verkman et al., 2017). Hence, changes in expression and subcellular distribution of AQP4 in astrocytes as found in the present study might lead to neurological complications, for instance an increased vulnerability to Alzheimer’s disease (Furman, Green, & Lane, 2023; Monllor et al., 2023; Silva et al., 2022), long after acute symptoms of COVID-19 have vanished. We did not find significant differences in brain weights between COVID-19 cases and controls, suggesting that impaired water homeostasis upon AQP4 dysregulation in COVID-19 does not evoke severe brain edema. It is not known whether AQP4 dysregulation lasts after acute SARS-CoV2 infection and more studies are needed to test long-term effects of COVID-19 on AQP4 polarization, cognitive abilities and mental health. Together with results of other studies on the role of astrocytes in COVID-19 and long-COVID (Huang & Fishell, 2022; Murta, Villarreal, & Ramos, 2020; Rosu et al., 2022; Steardo et al., 2022; Tremblay et al., 2020; Zorec & Verkhratsky, 2023), our results imply an important role for astrocytes in neuroinflammation as well as neurological and mental disorders due to SARS-CoV-2 infection.

## Acknowledgments

We thank A.C. Rakete for technical assistance.

