## Supplemental material for "Microgliosis, astrogliosis and loss of aquaporin-4 polarity in frontal cortex of COVID-19 patients"

### Supplementary information

Supplementary table 1: Clinical details of cases and controls

| Nr. | Age | sex | cause of death | Comorb. |
|---|---|---|---|---|
| Case #1 | 93 | m | Pneumonia | Septic multiorgan failure |
| Case #2 | 63 | m | Pulmonary embolism | Asthma, diabetes, adipositas |
| Case #3 | 52 | m | Pulmonary embolism | Hepato- and splenomegaly |
| Case #4 | 54 | f | Pneumonia | Adipositas |
| Case #5 | 56 | m | Pneumonia | Kidney disease, diabetes, adipositas |
| Case #6 | 84 | m | Pneumonia | Cardiac disease |
| Case #7 | 89 | f | Pneumonia | Hypertension |
| Case #8 | 81 | m | Pneumonia | Lung cancer |
| Case #9 | 83 | m | Pneumonia | Cardiac disease |
| Case #10 | 85 | f | Pneumonia | Cardiac disease, kidney disease |
| Case #11 | 59 | f | Pneumonia | Pulmonary embolism |
| Case #12 | 29 | m | Pneumonia | Septic multiorgan failure |
| Control #1 | 45 | m | Pericardial tamponade | Aortic dissection |
| Control #2 | 78 | m | Cardiac failure | Pancreatic cancer |
| Control #3 | 87 | f | Myocardial Infarction | Arteriosclerosis |
| Control #4 | 64 | m | Hemorrhagic shock | Cardiac disease, liver disease |
| Control #5 | 85 | m | Endocarditis | Liver cirrhosis, arteriosclerosis |
| Control #6 | 33 | f | Hemorrhagic shock | Aortic dissection |
| Control #7 | 51 | m | Myocardial infarction | Adipositas |
| Control #8 | 58 | m | Cardiac failure | Pneumonia |
| Control #9 | 92 | m | Cardiac failure | Cerebral infarction |
| Control #10 | 86 | f | Sepsis | Hypertension, diabetes |

Supplementary table 2: Primary / secondary antibody combinations

| Staining number | primary antibody | Host species | Source | Dilution | Antigen retrieval | Secondary antibody* | Dilution |
|---|---|---|---|---|---|---|---|
| 1 | Iba1 | rb | Wako 019-19741 | 1/100 | alkaline | Alexa488 anti rb | 1/500 |
| | CD31 | ms | Agilent Dako M082329 | 1/100 | | Alexa555 anti ms | 1/500 |
| 2 | S100B | rb | Agilent Dako Z0311 | 1/100 | acidic | Alexa488 anti rb | 1/500 |
| | GFAP | ch | Abcam ab4674 | 1/100 | | Alexa555 anti ch | 1/500 |
| 3 | GFAP | rb | Agilent Dako Z033429 | 1/200 | alkaline | Alexa488 anti rb | 1/500 |
| | CD31 | ms | Agilent Dako M082329 | 1/100 | | Alexa555 anti ms | 1/500 |
| 4 | AQP4 | rb | Sigma Hpa014784 | 1/100 | acidic | Alexa488 anti rb | 1/500 |
| 5 | GFAP | ch | Abcam ab4674 | 1/100 | alkaline | Alexa405 anti ch | 1/500 |
| | Cd31 | ms | Agilent Dako M082329 | 1/100 | | Alexa488 anti ms | 1/500 |
| | AQP4 | rb | Sigma Hpa014784 | 1/100 | | Alexa555 anti rb | 1/500 |

Mouse = ms, rabbit = rb, chicken = ch, *host species: goat
